# Neurogenesis-dependent transformation of hippocampal memory traces during systems consolidation

**DOI:** 10.1101/2025.06.05.658072

**Authors:** Ali Golbabaei, Cesar A.O. Coelho, Mitchell L. de Snoo, Sheena A. Josselyn, Paul W. Frankland

**Affiliations:** Program in Neuroscience & Mental Health, Hospital for Sick Children, Toronto, Canada; Institute for Medical Science, University of Toronto, Toronto, Canada; Department of Psychology, University of Toronto, Toronto, Canada; Department of Physiology, University of Toronto, Toronto, Canada; Canadian Institute for Advanced Research, Program in Child and Brain Development

**Keywords:** Episodic memory, systems consolidation, hippocampus, prelimbic cortex, adult neurogenesis, generalization, optogenetics, engram

## Abstract

Memories for events (i.e., episodic memories) change qualitatively with time. Systems consolidation theories posit that organizational changes accompany qualitative shifts in memory resolution, but differ as to the locus of this reorganization. Whereas some theories favor inter-regional changes in organization (e.g., hippocampus→cortex; multiple trace theory), others favor intra-regional reorganization (e.g., within-hippocampus; trace transformation theory). Using an engram tagging and manipulation approach in mice, here we provide evidence that intra-regional changes in organization underlie shifts in memory resolution. We establish that contextual fear memories lose resolution as a function of time, with mice exhibiting conditioned freezing in both the training apparatus (context A) and a second apparatus (context B) at more remote delays (freezing_A_ ≡ freezing_B_ at remote delay). By tagging either hippocampal (dentate gyrus) or cortical (prelimbic cortex) neuronal ensembles in context A, and then pairing their optogenetic activation with shock (in context C), we monitored the resolution of these artificially-generated memories at recent versus remote post-conditioning delays by testing mice in contexts A and B. Hippocampal engrams for a fear conditioning event were initially high-resolution (recent delay: freezing_A_ >> freezing_B_) but lost resolution with time (remote delay: freezing_A_ ≡ freezing_B_). In contrast, cortical engrams were initially low-resolution and remained low-resolution over time (recent and remote delay: freezing_A_ ≡ freezing_B_). Transformation of hippocampal engrams was dependent on adult hippocampal neurogenesis. Eliminating hippocampal neurogenesis arrested hippocampal engrams in a recent-like, high-resolution state where mice continued to exhibit discriminative freezing at remote delays.

**Graphical abstract:** 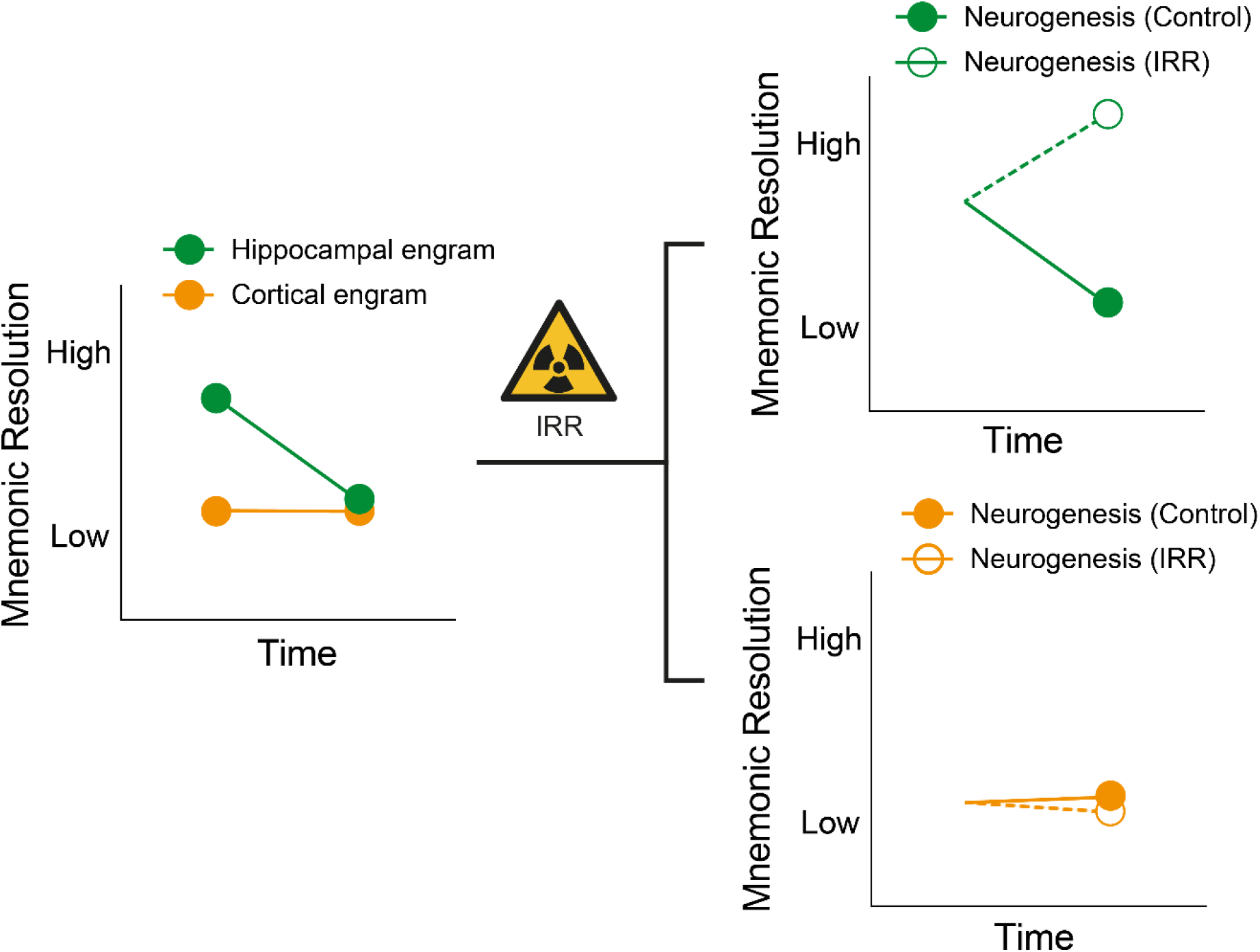

## Introduction

The mnemonic resolution of event-based (i.e., episodic) memories typically fades with time. For example, episodic memories may initially be highly resolved, where the memory contains event details and is bound to the specific spatial context in which the event occurred. However, with time, episodic memories may lose resolution, becoming less detailed, and no longer linked to a particular context^1^. Changes in memory organization are thought to accompany these shifts in mnemonic resolution, via a process known as systems consolidation^2,3^. One hypothesized benefit of systems consolidation-dependent changes in memory quality is that lower resolution episodic memories may not only guide behavior in situations that closely match encoding circumstances, but also in novel, but related, situations (i.e., promote generalization)^4–15^.

Models of systems consolidation differ as to whether inter-regional or intra-regional shifts in organization track changes in memory resolution. Multiple trace theory^16^ (MTT) proposes that inter-regional reorganization is responsible for time-dependent changes in memory resolution, assuming that the hippocampus encodes high resolution mnemonic information and the cortex low resolution information. MTT asserts that episodic memories become independent of the hippocampus once they lose mnemonic resolution. In contrast, trace transformation theory^17,18^ (TTT), and its derivatives^19^, proposes that intra-regional reorganization underlies changes in mnemonic resolution. This builds on human neuroimaging findings that show posterior to anterior shifts in hippocampal activation patterns accompany time-dependent changes in memory resolution, with remote, gist-like memories engaging the anterior pole of the hippocampus^20,21^.

Using an engram tagging approach in mice^22^, the goal of the current study was to determine whether intra- or inter-regional reorganization provides more accurate account of systems consolidation. In particular, we took advantage of a false memory paradigm developed by Ramirez and colleagues^23^ in order to isolate the contribution of hippocampal and cortical engrams to the expression of contextual fear conditioning. In that study, they used a TetTag system to express excitatory opsins in dentate gyrus (DG) ensembles activated by exposure to context A. Mice were subsequently fear conditioned (in the conditioning context, context C) while reactivating the context A ensembles. When later tested, mice froze in context A. Since context A was never paired with shock, this indicates that the mice formed a ‘faIse’ context A-shock association. Moreover, this false memory exhibited specificity. Mice did not freeze when they were tested in a second context, context B, suggesting that DG engrams support high resolution event-based memories, at least soon after conditioning.

Here we adapted this approach and paired activation of either hippocampal (DG) or cortical (prelimbic cortex; PrL^24^) engram ensembles (tagged in context A) with conditioning (in context C). By subsequently evaluating freezing in contexts A and B either 2- or 14-days following conditioning, we were able to track how the mnemonic resolution of hippocampal and cortical engrams changes with the passage of time. Consistent with intra-regional accounts of systems consolidation (e.g., TTT), we find that hippocampal engrams are initially high resolution, but lose resolution over time. In contrast, cortical engrams are initially low-resolution, and remain low-resolution at more remote delays. We further establish that high→low shifts in memory resolution of hippocampal engrams are modulated by hippocampal neurogenesis.

## Results

### Contextual fear memories lose precision with time

The binding of events to their surrounding spatial context is a core feature of episodic memory and may be modeled in rodents using contextual fear conditioning^25^. To explore how the resolution of contextual fear memories changes with time, we trained mice and then tested the specificity of their memory by placing them in the original training context (context A) or a second context (context B) at recent (2 days) or remote (14 days) post-training delays and measured freezing, a species-typical fear behavior^26^ (Fig. 1A). As in previous studies^27–29^, in these tests of context-specificity we operationally defined discriminative freezing (freezing_A_ >> freezing_B_) as reflecting high mnemonic resolution and non-discriminative freezing (freezing_A_ ≡ freezing_B_) as reflecting low mnemonic resolution^27–33^. At the recent delay, mice exhibited high-resolution contextual fear memories, freezing more in context A compared to context B. In contrast, when mice were tested 14 days following training, they exhibited elevated freezing in context B (Fig. 1B), suggesting that memory for the conditioning event loses resolution with time. This loss of mnemonic resolution is supported by delay-dependent reductions in discrimination between contexts A and B (Fig. 1C).

**Figure 1.**
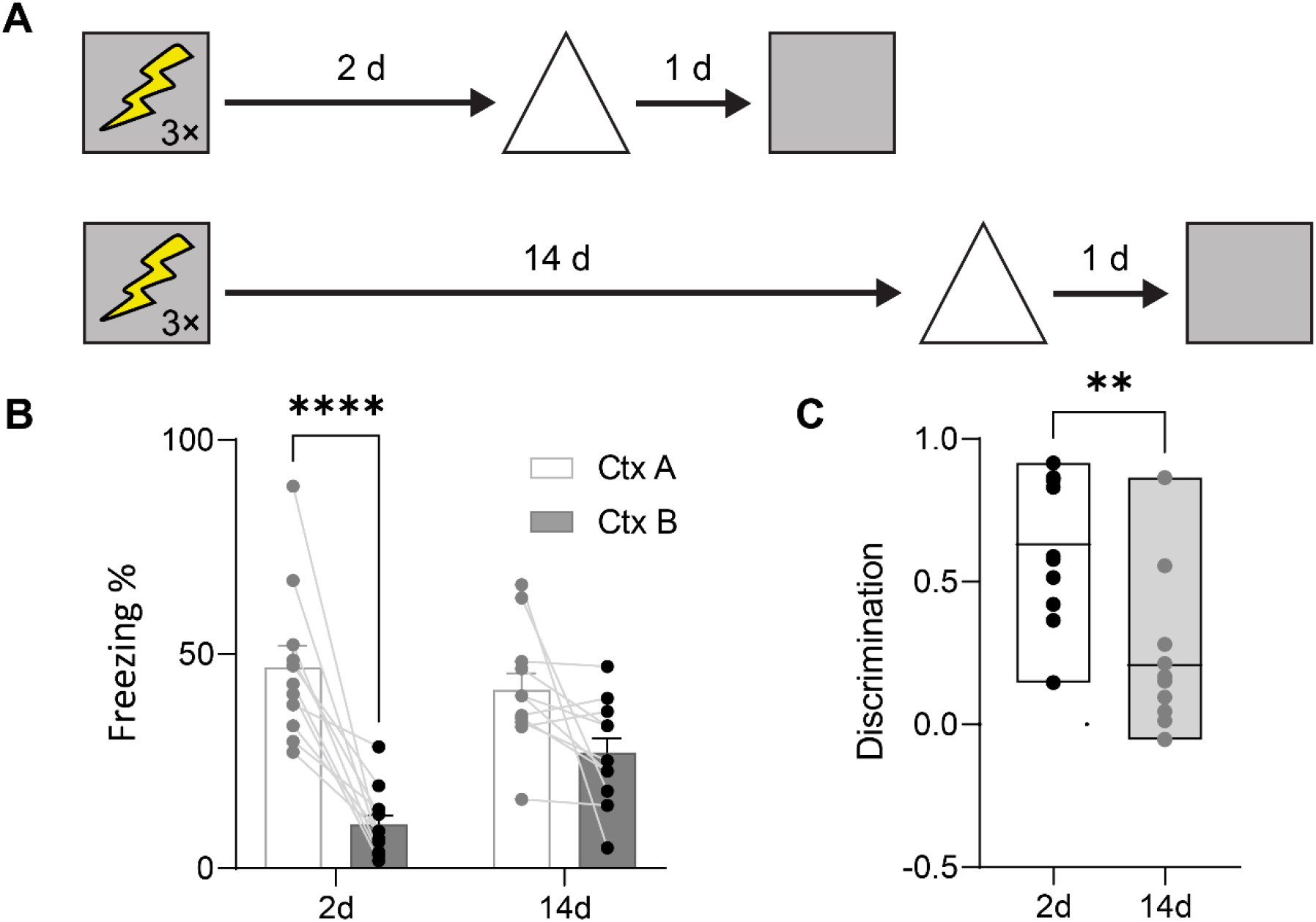
Contextual memories lose resolution with time. (A) Experimental design. Mice were trained in contextual fear conditioning and tested either 2 or 14 d later in the conditioned context (square) or a similar context (triangle). (B) Mice froze more in the conditioned context compared to the similar context at the recent delay but not the remote delay (2-way ANOVA, main context effect: *F*_1,20_ = 32.82, *P* < 0.001; main delay effect: *F*_1,20_ = 2.46, *P* = 0.13; context × delay interaction: *F*_1,20_ = 6.04, *P* = 0.02). (C) Context discrimination declined with retention delay (unpaired t-test, t_20_ = 3.72, *P* < 0.01). * *P* ≤ 0.05, ** *P* ≤ 0.01, *** *P* ≤ 0.001, **** *P* ≤ 0.0001.

These results replicate earlier studies that show time-dependent decreases in mnemonic resolution of contextual fear memories in mice^27,28,30^ and rats^29^.

### Hippocampal-based contextual engrams lose resolution over time

To evaluate whether the mnemonic resolution of hippocampal engrams changes with time, we microinjected an adeno-associated virus (AAV) expressing Robust Activity Marking (RAM)-ChR2 into the DG. Temporary removal of doxycycline from their diet allowed us to tag DG ensembles active with ChR2 when the mice were exposed to context A. Two days later mice were placed in the conditioning context (context C) while bluelight (BL) photostimulation was applied to reactivate ensembles tagged in context A. Mice were subsequently tested in context A and a second context, context B, either 2 or 14 days later (Fig. 2A), allowing us to track how the mnemonic resolution of this DG-based engram changes with time. At the recent delay mice froze more in context A compared to context B, indicating that the DG engram supported expression of a high-resolution memory that showed context-specificity at a recent delay, similar to previous studies^23^. In contrast, at the remote delay, mice exhibited elevated freezing in context B, suggesting that memory resolution of the tagged context A engram decreases with time (Fig. 2B). Consistent with this interpretation, we found that discrimination decreased as a function of memory age (Fig. 2C).

**Figure 2.**
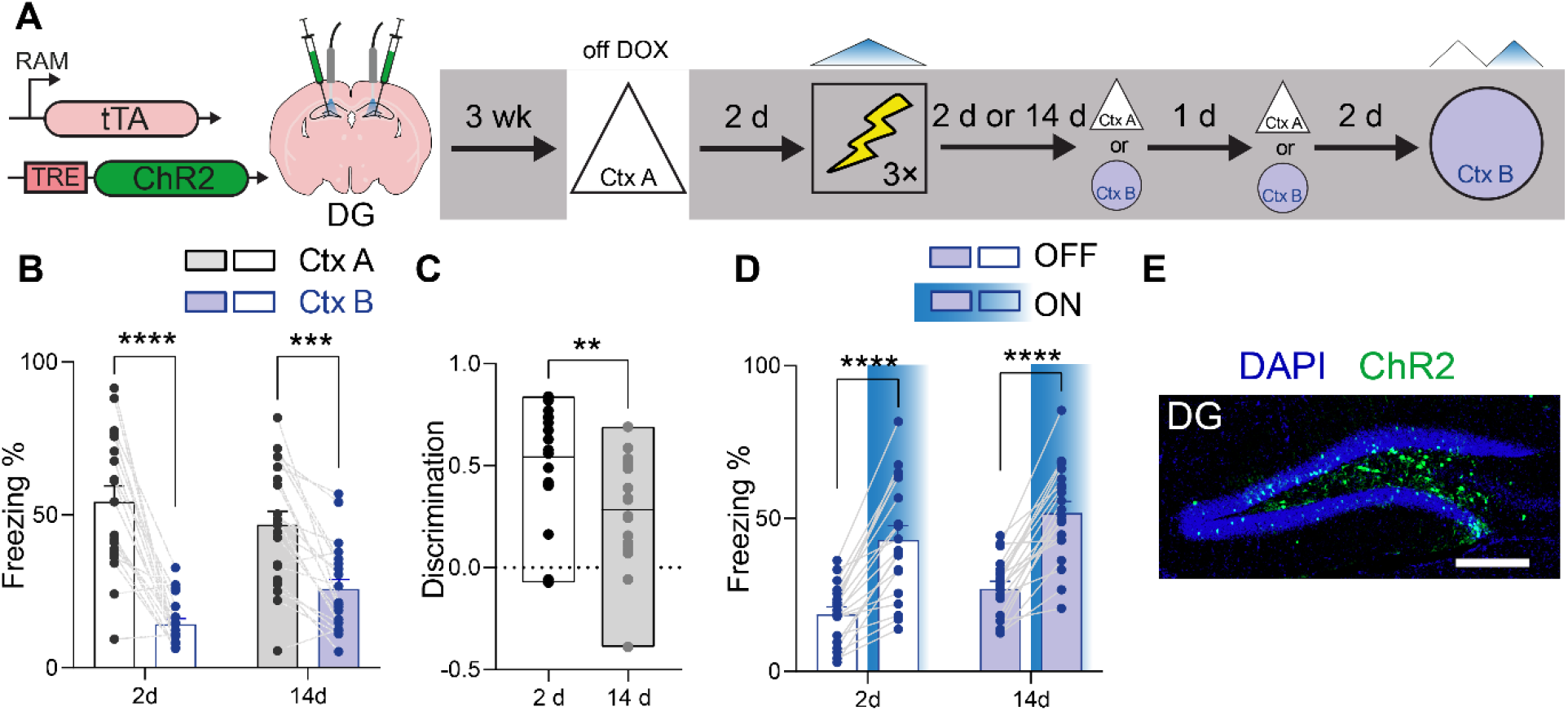
DG engrams lose resolution at remote delays. (A) Mice were microinjected with AAV expressing RAM-ChR2 in DG and were exposed to context A while off DOX to tag active neuronal ensembles. Two days later, photostimulation of the tagged ensemble was paired with shock in context C and mice were tested in either context A or B at either 2 or 14 d post-conditioning. Two days following the completion of these tests, mice were retested in context B in the presence/absence of BL photostimulation. (B) Freezing in contexts A and B at recent and remote delays (2-way ANOVA, main context effect: *F*_1,37_ = 83.37, *P* < 0.001; main delay effect: *F*_1,37_ = 0.20, *P* = 0.65; context × delay interaction: *F*_1,37_ = 8.09, *P* < 0.01). (C) Reduced discrimination at the remote delay (unpaired t-test, t_37_ = 3.04, *P* < 0.01). (D) BL photostimulation of tagged DG neurons increased freezing in context B at recent and remote delays (2-way ANOVA, main stimulation effect: *F*_1,37_ = 137.8, *P* < 0.001; main delay effect: *F*_1,37_ =4.58, *P* = 0.04; stimulation × delay interaction: *F*_1,37_ = 0.01, *P* = 0.91). (E) Representative image showing ChR2-tagged (green) context A engram neurons in DG. Scale bar = 250 µm. * *P* ≤ 0.05, ** *P* ≤ 0.01, *** *P* ≤ 0.001, **** *P* ≤ 0.0001.

Effective ChR2-tagging of DG engrams was subsequently confirmed by testing mice in context B, with and without BL photostimulation (Fig. 2A). BL photostimulation induced freezing (Fig. 2D), consistent with ChR2-expressing neurons being confined to granule cell layer of DG (Fig. 2E). When conditioning occurred without reactivation of DG engram ensembles (tagged in context A), mice exhibited no freezing when subsequently tested in contexts A or B either 2 or 14 days later (Fig. S1).

This indicates that exposure to shock alone in the conditioning context (in the absence of BL photostimulation) does not induce generalized freezing in either contexts A or B.

### Cortical-based contextual engrams are low-resolution, but do not change over time

We next tested whether the mnemonic resolution of cortical engrams differs from hippocampal engrams, and whether their resolution changes with time. To do this, we microinjected AAV-RAM-ChR2 into prelimbic cortex (PrL), a frontal cortical region that is important for consolidation of contextual fear memory^24,34–36^. Removal of DOX upon exposure to context A allowed us to tag context A PrL ensembles with ChR2, and subsequently reactivate these ensembles with BL as mice were conditioned two days later (Fig. 3A). In subsequent retrieval tests, we found that mice exhibited elevated levels of freezing in context B at both recent and remote delays (Fig. 3B). This suggests that cortical engrams support low-resolution memories of the conditioning event, and resolution does not change significantly as a function of retention delay (Fig. 3C). Supporting this interpretation, analysis of discrimination scores indicated that they were a) lower than for hippocampal engrams at the recent delay, and b) remained low at the remote delay (Fig. S2). Effective ChR2-tagging of PrL engrams was subsequently confirmed by testing mice in context B, with and without BL photostimulation (Fig. 3A). BL photostimulation induced freezing (Fig. 3D), consistent with robust ChR2-labeling of engram neurons throughout the PrL (Fig. 3E).

**Figure 3.**
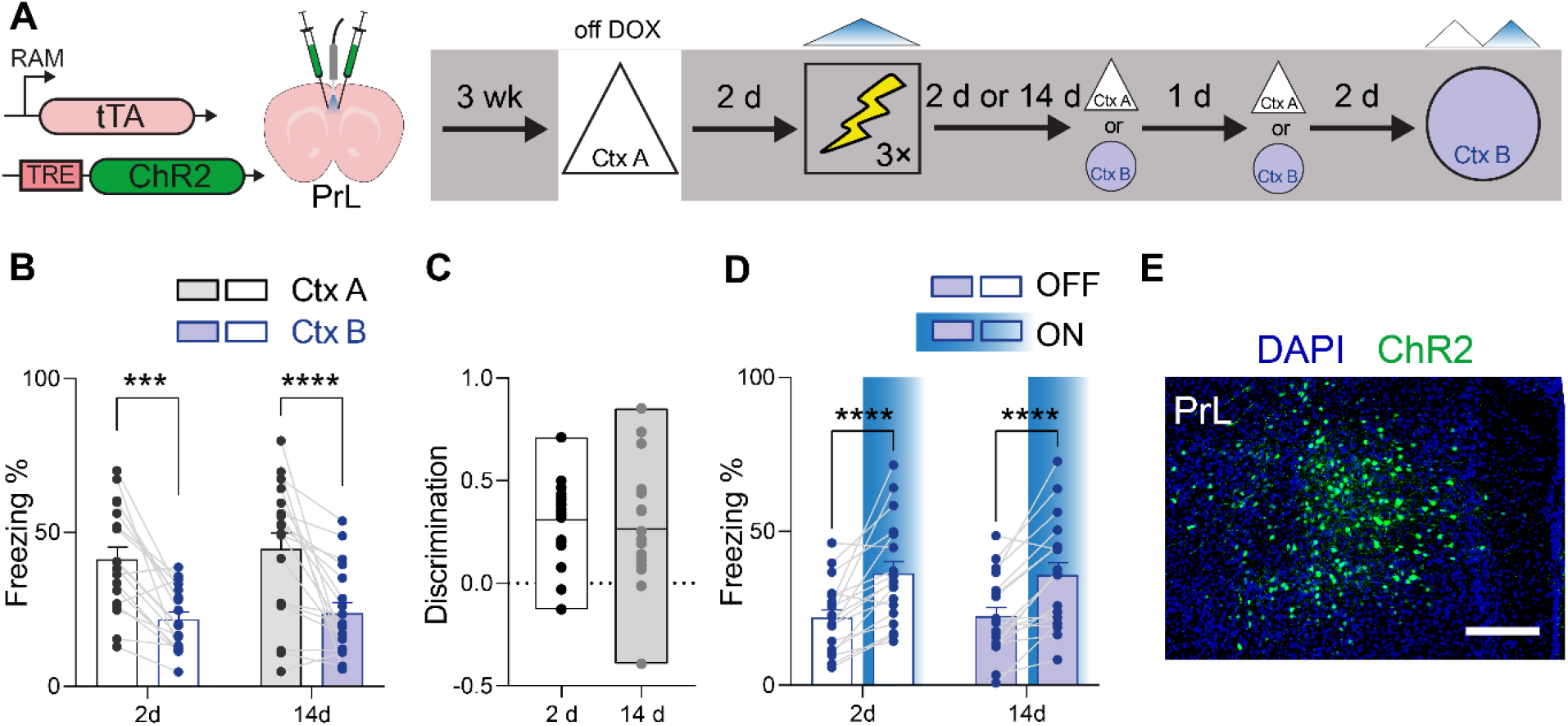
PrL engrams are low resolution at recent and remote delays. (A) Mice were microinjected with AAV expressing RAM-ChR2 in PrL and were exposed to context A while off DOX to tag active neuronal ensembles. Two days later, photostimulation of the tagged ensemble was paired with shock in context C and mice were tested in either context A or B at either 2 or 14 d post-conditioning. Two days following the completion of these tests, mice were retested in context B in the presence/absence of BL photostimulation. (B) Freezing in contexts A and B at recent and remote delays (2-way ANOVA, main context effect: *F*_1,34_ = 52.96, *P* < 0.001; main delay effect: *F*_1,37_ = 0.35, *P* = 0.56; context × delay interaction: *F*_1,34_ = 0.05, *P* = 0.82). (C) Discrimination does not change with time (unpaired t-test, t_34_ = 0.54, *P* = 0.59). (D) BL photostimulation of tagged PrL neurons increased freezing in context B at recent and remote delays (2-way ANOVA, main stimulation effect: *F*_1,34_ = 48.91, *P* < 0.001; main delay effect: *F*_1,37_ = 0.00, *P* = 0.99; stimulation × delay interaction: *F*_1,34_ = 0.06, *P* = 0.81). (E) Representative image showing ChR2-tagged (green) context A engram neurons in PrL. Scale bar = 250 µm. * *P* ≤ 0.05, ** *P* ≤ 0.01, *** *P* ≤ 0.001, **** *P* ≤ 0.0001.

### Eliminating hippocampal neurogenesis prevents high→low shifts in memory resolution of hippocampal engrams

One factor that is hypothesized to transform hippocampal engrams is ongoing neurogenesis^30,37–40^.

For example, we recently found that neurogenesis-induced remodeling of hippocampal engram circuits supporting a contextual fear memory enables CA3 and CA1 engram neurons to be promiscuously active in a broader set of situations beyond the original training context (i.e., engram neurons that are usually selectively active in context A become active in context B)^30^. Based on this, we hypothesized that eliminating hippocampal neurogenesis might ‘arrest’ hippocampal engram circuits in a recent memory state, where hippocampal engrams support high-resolution contextual fear memories, even when tested at remote delays.

To test this hypothesis, we combined the cortical (PrL) and hippocampal (DG) engram tagging and manipulation strategies outlined above with irradiation (IRR)-induced ablation of hippocampal neurogenesis^41^. Mice received a single dose of focal gamma IRR (or SHAM treatment) two weeks prior to DOX removal and exposure to context A. Mice were returned to the DOX diet to close the tagging window and then two days later conditioned in context C, during which photostimulation of the context A tagged ensembles was paired with shock. Conditioned fear was subsequently evaluated in contexts A and B 14 days later (Fig. 4A).

**Figure 4.**
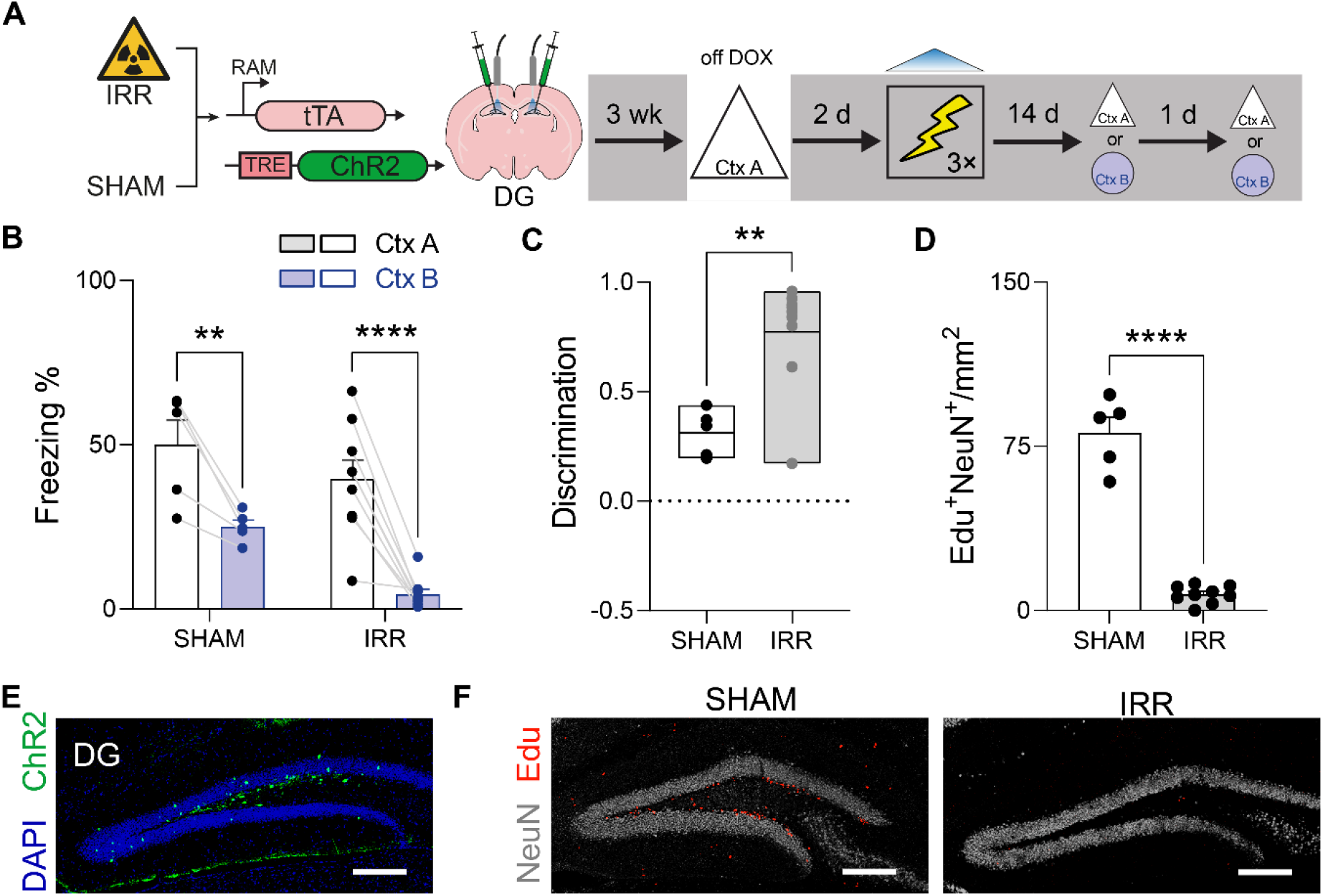
Elimination of neurogenesis prevents loss of DG engram resolution. (A) IRR or SHAM-treated mice were microinjected with AAV expressing RAM-ChR2 in DG and placed in context A while off DOX to tag the active ensembles. Two days later, photostimulation of the tagged ensemble was paired with shock in context C and mice were tested in either context A or B at either 2 or 14 d post-conditioning. (B) Freezing in contexts A and B at recent and remote delays (2-way ANOVA, main context effect: *F*_1,12_ = 53.55, *P* < 0.001; main irradiation effect: *F*_1,12_ = 7.48, *P* = 0.02; context × irradiation interaction: *F*_1,12_ = 1.54, *P* = 0.24). (C) Elevated discrimination in IRR compared to SHAM mice (unpaired t-test, t_12_ = 3.93, *P* < 0.01). (D) IRR reduced DG neurogenesis (i.e., EdU^+^/NeuN^+^ cells) (unpaired t-test, t_12_ = 13.39, *P* < 0.001). (E) Representative image showing ChR2-tagged (green) context A engram neurons in DG. Scale bar = 250 µm. (F) Representative Edu labeling (red) in DG in SHAM (left) vs. IRR (right) mice. Scale bar = 250 µm. * *P* ≤ 0.05, ** *P* ≤ 0.01, *** *P* ≤ 0.001, **** *P* ≤ 0.0001.

SHAM-treated mice in which reactivation of context A-tagged DG engram neurons was paired with shock exhibited high levels of freezing in context B when tested 14 days post-conditioning, consistent with our previous experiment showing that DG engrams lose mnemonic resolution as a function of time. However, IRR prevented this time-dependent loss of mnemonic resolution. IRR-treated mice in which reactivation of context A-tagged DG engram neurons was paired with shock exhibited context specificity, freezing in context A but not in context B when tested at the remote delay (Fig. 4B). Consistent with this interpretation, discrimination scores were elevated in IRR, compared to SHAM, mice (Fig. 4C). Histological analyses confirmed that IRR reduced hippocampal neurogenesis by approximately 80%, without affecting engram labeling in DG (Fig. 4D-F). These results support our hypothesis that elimination of hippocampal neurogenesis prevents transformation of hippocampal engrams, crystallizing them in a recent-like memory high-resolution state even at remote delays.

Mice in which reactivation of context A-tagged PrL engram neurons was paired with shock (Fig. 5A) exhibited high levels of freezing in context A and elevated levels of freezing in context B, indicating that PrL engrams maintain low-resolution contextual fear memories. Freezing in context B was equivalently elevated in SHAM and IRR groups, indicating that IRR treatment does not impact the mnemonic resolution of PrL engrams (Fig. 5B-C). Histological analyses confirmed that IRR reduced hippocampal neurogenesis by approximately 80%, without affecting engram labeling in PrL (Fig. 5D-F). Therefore, these results indicate that elimination of hippocampal neurogenesis does not impact the fate of PrL-based contextual fear engrams, which remain in a low-resolution state at remote delays.

**Figure 5.**
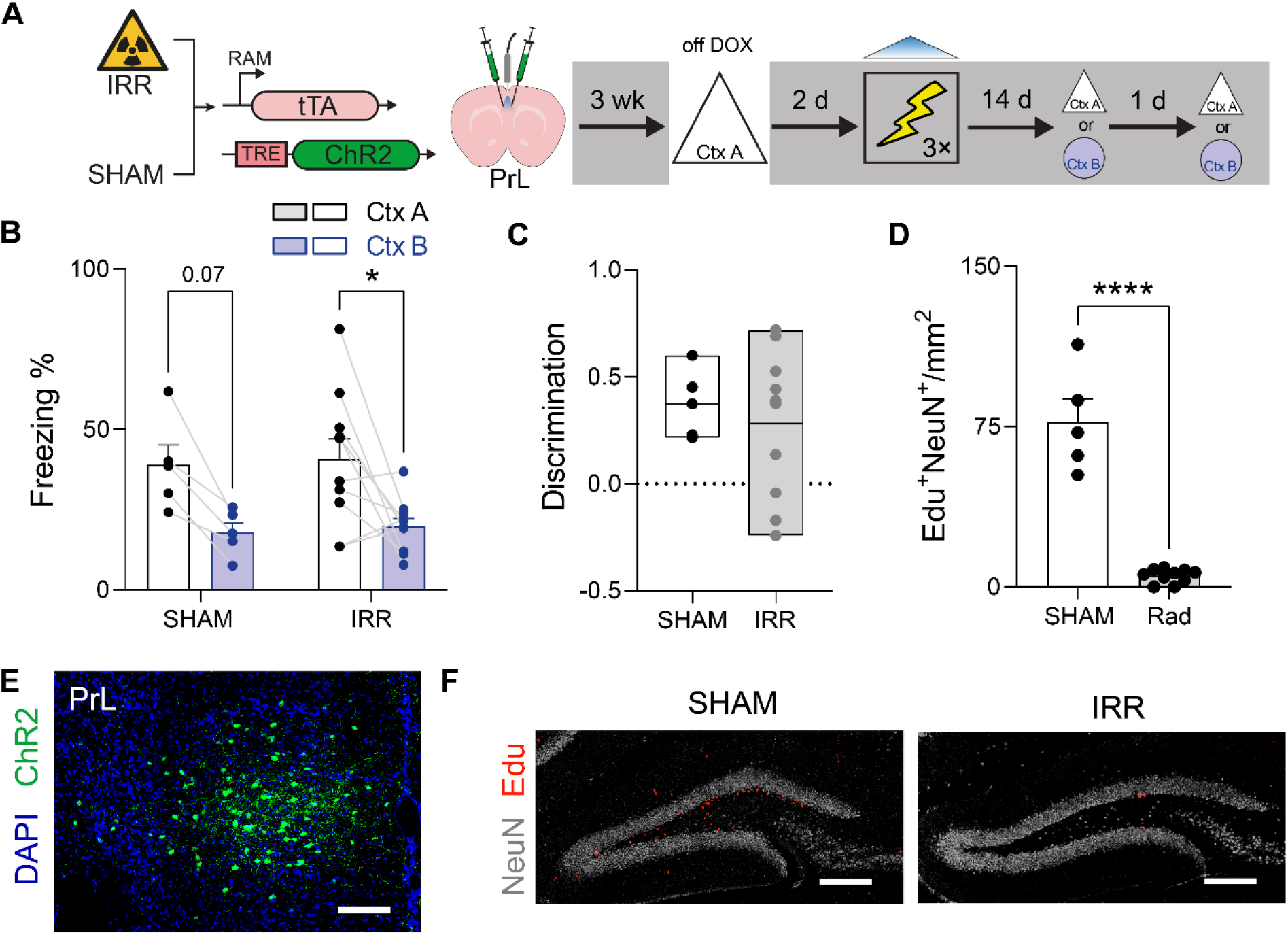
Elimination of neurogenesis does not affect PrL engram resolution. (A) IRR or SHAM-treated mice were microinjected with AAV expressing RAM-ChR2 in PrL and placed in context A while off DOX to tag the active ensembles. Two days later, photostimulation of the tagged ensemble was paired with shock in context C and mice were tested in either context A or B at either 2 or 14 d post-conditioning. (B) Freezing in contexts A and B at recent and remote delays (2-way ANOVA, main context effect: *F*_1,13_ = 14.03, *P* < 0.01; main irradiation effect: *F*_1,13_ = 0.10, *P* = 0.75; context × delay interaction: *F*_1,13_ = 0.00, *P* = 0.98). (C) Similar discrimination in SHAM vs. IRR mice (unpaired t-test, t_13_ = 0.56, *P* = 0.59). (D) IRR reduced DG neurogenesis (i.e., EdU^+^/NeuN^+^ cells) (unpaired t-test, t_12_ = 13.39, *P* < 0.001). (E) Representative image showing ChR2-tagged (green) context A engram neurons in PrL. Scale bar = 250 µm. (F) Representative Edu labeling (red) in DG in SHAM (left) vs. IRR (right) mice. Scale bar = 250 µm. * *P* ≤ 0.05, ** *P* ≤ 0.01, *** *P* ≤ 0.001, **** *P* ≤ 0.0001.

## Discussion

In this study we used an engram false memory procedure^23^ to isolate hippocampal (DG) and cortical (PrL) engrams and study how their mnemonic resolution changes as a function of time. Our experiments suggest three main conclusions. First, hippocampal engrams for a fear conditioning event are initially high-resolution but lose resolution with time. Second, cortical engrams for a fear conditioning event are initially low-resolution and remain low-resolution over time. Third, the high-resolution → low-resolution transformation of hippocampal engrams is modulated by hippocampal neurogenesis. Eliminating hippocampal neurogenesis, by delivering focal gamma irradiation, arrested contextual fear memories in a recent-like state where mice continued to exhibit high-resolution memories (i.e., discriminative freezing) at remote post-conditioning delays.

Systems consolidation theories propose that shifts in memory resolution are accompanied by organizational changes at the level of engrams, with some theories favoring inter-regional changes in organization (e.g., hippocampus→cortex; MTT^16^) and others favoring intra-regional changes in organization (e.g., within-hippocampus; TTT^17,18^). The current pattern of results is more compatible with the latter; that is, engrams corresponding to contextual fear memories are transformed within the hippocampus. Such an enduring role for the hippocampus in event-linked memories is consistent with recent behavioral studies in rodents and neuroimaging and neuropsychological studies in humans. In rodents, lesion and inactivation studies indicate that the hippocampus is important for the expression of event-linked memory (including contextual fear memories) at remote delays^42–44^. In humans, the hippocampus remains engaged during retrieval of remote, event-linked, gist memories at remote time points^20,21^.

What drives transformation of event-linked memories within the hippocampus? We found that focal gamma irradiation arrested DG-based (but not PrL-based) contextual fear engrams in a high-resolution state that is typically only observed at recent post-conditioning delays. This indicates that ongoing hippocampal neurogenesis is necessary for the transformation of hippocampal memories, supporting earlier proposals that hippocampal neurogenesis modulates the progression of systems consolidation^37–39^. For example, Kitamura et al^38^ found that focal irradiation eliminated hippocampal neurogenesis, and maintained contextual fear memories in a state where they continued to be sensitive to pharmacological inactivation of the dorsal hippocampus even at remote post-conditioning delays. Since adult hippocampal neurogenesis is regulated by many variables^45^, this raises the testable hypothesis that factors such as age and environment modulate the rate of intra-hippocampal memory transformation.

The precise mechanistic basis for memory transformation was not studied here. However, our recent work suggests that transformation involves neurogenesis-dependent rewiring of corresponding hippocampal engrams^30^. In that study, we tracked the connectivity of hippocampal engrams supporting contextual fear memory over time using e-GRASP^46^ and identified delay-dependent changes in hippocampal engram connectivity. These included two types of wiring changes that would serve to increase the likelihood of CA3 and CA1 engram neuron activation: a) delay-dependent decreases in DG engram neuron filopodial contacts onto CA3 PV^+^ interneurons^32^ and b) delay-dependent increases in excitatory CA3→CA1 engram-specific connectivity^46^.

Consistent with this, we found that CA3 and CA1 engram neurons were promiscuously activated in related situations (i.e., context B) that did not match the original training conditions (i.e., context A), and their promiscuous activation predicted with the emergence of event-linked gist memory (reflected by elevated fear in context B). Importantly, we found that newborn neurons directly contact CA3 engram neurons, and eliminating neurogenesis prevented the delay-dependent reorganization of hippocampal engram circuits and the emergence of event-linked gist memory. Conversely, promoting neurogenesis accelerated reorganization of hippocampal engram circuits, leading to earlier emergence of event-linked gist memory^30^.

Neurogenesis suppression following focal irradiation is nearly complete and therefore focal irradiation serves as a powerful intervention to address the hypothesis that hippocampal neurogenesis is necessary for delay-dependent transformation of hippocampal engrams. Nonetheless, focal irradiation has off-target effects (including neuroinflammation and illness^41,47^), and altered memory function may be related to these effects rather than the elimination of adult neurogenesis. However, several observations suggest that the abnormal maintenance of high-resolution contextual fear memories following irradiation is linked to disruptions of neurogenesis, rather than off-target effects. First, preserved high-resolution context memories were only observed for DG-, but not PrL-based, engrams. The absence of effects on PrL-based engrams suggests that irradiation does not nonspecifically alter expression of fear behaviors in mice. Moreover, since preservation of high-resolution contextual fear memory may be considered as a gain-of-function memory phenotype, it seems unlikely that preserved discriminatory freezing is related to either neuroinflammatory responses or illness. Second, when more selective methods are used to manipulate hippocampal neurogenesis in adult mice^48^, contextual and spatial memories are similarly maintained in high-resolution states at remote delays^30^.

Preserved contextual discrimination following irradiation contrasts with a large number of studies showing that hippocampal neurogenesis contributes to behavioral pattern separation^49,50^. In many of these studies, behavioral pattern separation was assessed using contextual fear conditioning protocols similar to those used here. However, where our approach differs crucially is in the timing of irradiation with respect to training. In our experiment, irradiation occurred 2 weeks before training and memory was assessed at a remote (as well as recent) delay post-conditioning. Because new neurons do not become synaptically integrated into hippocampal circuits until at least ~16 days^51^, this ensures that any effects on remote memory are most likely related to the absence of integrating new neurons (~2-4 weeks in age) during the post-conditioning window. In contrast, in previous studies irradiation typically occurred several weeks prior to training, and memory was tested at recent post-conditioning delays. Such protocols assess, therefore, the impact of reduced neurogenesis at the time of training on memory formation.

In the current study, a number of limitations are worth noting. First, in these experiments we paired hippocampal (DG) or cortical (PrL) engram activation with shock during conditioning. This allowed us to track resolution changes in engrams that were either predominantly DG- or PrL-based.

However, it is highly unlikely that activation was limited to the targeted engram during conditioning (e.g., when DG engram activation was paired with shock, it is likely that PrL-based circuits were also engaged). Nevertheless, this type of contamination associated with activation of non-engram substrates appears to be minimal given that we observed distinct trajectories of mnemonic resolution depending on whether we targeted DG- vs. PrL-based engrams. Second, in the RAM-based engram-tagging system, transgene expression declines beyond ~2 weeks and, therefore, we assessed remote memory only 14 days post-conditioning. Systems consolidation likely extends beyond this 2-week window in rodents^52,53^, and therefore future studies should use more enduring tagging systems that allow memory assessment at longer post-conditioning delays where loss of mnemonic resolution may be even more pronounced. Third, to address our hypothesis we relied solely on contextual fear conditioning. While a similar pattern of results has been observed using another aversively-motivated task (water maze^30^), future studies should explore whether these results generalize to other hippocampus-dependent tasks, especially using appetitively-motivated paradigms.

## Acknowledgments

We thank members of the JF lab for their input and encouragement at various stages of this project. This work was funded by grants from the National Institutes of Health (NIH) (R01 MH119421 to PWF and SAJ), Brain Canada (to PWF and SAJ), and the Canadian Institutes of Health Research (CIHR) (PJT180530 to PWF).

## Author Contributions

A.G., C.A.O.C., S.A.J. and P.W.F. conceived the project and designed the experiments. A.G. and C.A.O.C. conducted the behavioral experiments and histological analyses. M.d-S. conducted irradiation. A.G. and P.W.F. wrote the paper.

## METHODS

### Mice

In these experiments, we used wild-type (WT) mice. WT mice were generated ‘in-house’ by crossing C57BL/6NTac × 129S6/SvEvTac and the F1 descendants were used for all experiments. Mice were bred and maintained in the vivarium at the Hospital for Sick Children. They were maintained on a 12-hour light/12-hour dark cycle at all times with lights on between 8 AM to 8 PM each day. Food and water were provided ad libitum. The litters, usually containing 4-12 pups, were weaned 21 days post date of birth and housed 2-5 mice per cage. Males and females were housed separately and were used in all experiments. Eight to ten-week-old mice were used for behavioural experiments, which occurred during the light phase. All procedures were approved by the Animal Care Committee at the Hospital for Sick Children and Use Committee policies and confirmed the guidelines of the Canadian Council on Animal Care and the National Institute of Health on the Care and Use of Laboratory Animals.

### Contextual fear conditioning apparatus

#### Context configurations

For the generalization experiment (Fig. 1), the conditioning context (square in Fig. 1) had clear acrylic walls on the front and top of the chamber and aluminum walls on the sides and back of the chamber (32 × 25 × 25 cm). The floor consisted of metal grids that were used for administering foot shocks (0.7 mA, 2 s duration). A camera was installed in front of the acrylic wall for monitoring behavior. The similar context (triangle in Fig. 1) was located in a separate experimental room. This context had clear acrylic walls on the front and top of the chamber and aluminum walls on the sides and back (31 cm x 24 cm x 21 cm). The floor consisted of a white plastic flat plate.

For the inception experiments (Figs. 2–5), three different conditioning apparatuses were used (contexts A, B and C). In Context A, the side and back walls were covered with a C-shaped white plastic insert, and the floor covered with a flat white plastic insert (31 cm × 24 cm × 21 cm; Med Associates, St. Albans, VT). The illumination source was the room light. Cameras above the contexts recorded behavior. Context B consisted of a transparent plastic home cage in the center of a white chamber (50 × 18 × 45 cm). A camera mounted above recorded behavior, and the room was illuminated by a dim light. The conditioning context, context C (32 × 25 × 25 cm, Med Associates, St. Albans, VT), consisted of a front, top and back made of transparent acrylic pieces, two sidewalls made of modular aluminum and stainless-steel rods through which the foot shocks were delivered. The context was housed within a sound-attenuating chamber with fans (60 dB) providing background noise. During laser-stimulation and shock pairing experiments no illumination was provided in the chamber to reduce contextual information. Front-mounted cameras recorded behavior in near infrared light.

### Contextual fear conditioning procedures

#### Context tagging

For 3 consecutive days prior to starting the experiments, the fiber implants were connected to the optic cable for 2-5 mins in order to habituate mice to the procedure. Next, mice were removed from DOX diet (40 mg/kg) for 48 h, and placed in context A for 10 mins to tag neurons active in this context. After tagging, mice were put back in their home cage on DOX diet to close the tagging window.

#### Conditioning

On the day of the conditioning, mice were taken from the vivarium and temporarily held in a quiet holding area for 60 mins. They were transferred to the conditioning room and placed into the conditioning chamber (context C). Shocks were delivered at 120 s, 150 s, and 180 s after placement in the context and mice remained in the chamber for an extra 60 s after the final shock. They were returned to the holding room for 60-120 mins, and then transferred back to the vivarium. For the inception experiments, mice were connected to the optical fibers before placement in the conditioning context (context C). During the conditioning session, photostimulation (473 nm, 1-2 mW, 15 ms pulse duration) was delivered continuously at a frequency of 4 Hz for PrL tagging and 20 Hz for DG tagging, respectively.

#### Retrieval tests in contexts A and B

Mice were brought back to the holding room 60 mins prior to testing either 2 d or 14 d following conditioning. They were tested on consecutive days in contexts A and B (order counter-balanced, test duration 4 mins). In these tests, mouse behavior was monitored via an overhead camera (3.75 frames/s), and freezing was automatically tracked using FreezeFrame software. The percentage of time spent freezing time across the entire testing period was used for statistical analyses.

#### Optogenetic engram reactivation (in context B)

One day following testing in contexts A and B, mice were additionally tested in context B for 8 mins. Photostimulation was delivered for the second 4 mins of the test, and freezing was scored manually (because the presence of optical cables precluded automated scoring).

### Viral vectors

To tag active neuronal ensembles, we used viral vectors with the pAAV-RAM-d2TTA::TRE-ChR2-WPREpA plasmid (#84471, Addgene, Cambridge, MA) synthetized under DJ serotype in-house. The AAVs were produced in HEK293T cells with the AAV-DJ Helper Packaging System (Cell Biolabs, Inc., cat# VPK-400-DJ), according to the protocol of the manufacturer. Viral particles were filtered using the Virabind AAV Purification Kit (Cell Biolabs, Inc., cat#VPK-140). The final titer of the virus was set to approximately 1.0 ×10^11^ units/ml. The RAM system has the RAM promoter incorporated in the Tet-Off system, which allowed us to control ChR2 expression by withdrawal of Doxycycline from the diet (Off DOX^54^). When Off DOX, the RAM promoter expression is driven by expression of either Fos or Npas4 proteins, providing a way to robustly tag active neurons^54^. As a control, we used a version of this viral vector that expressed GFP instead of ChR2-eYFP.

### Stereotaxic surgery

#### Virus microinfusion

Before surgery mice were treated with atropine sulfate (0.1 mg/kg) and chloral hydrate (400 mg/kg) via intraperitoneal (i.p.) injection. Once a stable level of anesthesia occurred, mice received a subcutaneous (s.c.) injection of Meloxicam (4 mg/kg) for post-surgery analgesia.

The top of the scalp was disinfected with ethanol (70%), the scalp incised and retracted. A hole was made above the region of interest using an electric hand drill and a glass micropipette was inserted into the region for injection. The glass micropipette was filled with the virus solution and was connected to a Hamilton microsyringe via polyethylene tubing. A syringe pump was used to infuse the virus solution at a rate of 0.1 µl per minute. To target regions of interest the following coordinates were used (with respect to bregma): DG (AP −2.2 mm, ML ±1.5 mm, DV −2.3 mm) and PrL (AP +1.7 mm, ML ±0.35 mm, DV −1.8 mm). After infusion of the virus, the micropipette was held in place for an extra 10 mins (to maximize diffusion of the solution into the tissue) and then slowly retracted and the incision was sutured. Next, an optical fiber was implanted for delivery of photostimulation. The optical implant was constructed by attaching a fiber (200 µm diameter, 0.37 numerical aperture, Thorlabs) to a ceramic zirconia ferrule (1.25 mm diameter, Thorlabs) with epoxy resin. For PrL a single fiber was implanted above the injection sites at the midline, allowing delivery of photostimulation to both hemispheres. For DG two fibers were implanted bilaterally above injection sites on either side of the brain. Implants were secured on the skull by the application of black dental cement and screws. The wound was treated with Polysporin and the mice received a 1 ml injection of saline (0.9%, s.c.) for hydration. During recovery, mice were placed on a clean surface with access to a heating pad. After recovery, the mice were transferred to a clean cage and returned to the vivarium. The health of the mice was monitored for 3 days post-surgery. The behavioral procedures took place approximately 3 weeks after surgery.

### IRR-induced ablation of adult neurogenesis

#### Ablating newborn neurons

Focal gamma irradiation was used to eliminate adult neurogenesis^41^. Mice were anaesthetized and received a single dose of gamma irradiation (10 Gy) to the head, with the rest of the mouse body shielded by a lead chamber. This dose ablates neural progenitor cells by eliminating their proliferative capacity as well as shifting their progenitor cell fate toward a glial fate^41^. This results in an irreversible reduction in adult hippocampal neurogenesis in the DG across all phases of the experiment.

#### Quantification of adult neurogenesis

To measure the level of neurogenesis, 5-ethynyl-2’-deoxyuridine (EdU), a thymidine analog, was used. Edu integrates into the DNA of newly dividing cells^55^ allowing quantification of newly born cells. EdU (10 mg) was suspended in 1 ml of PBS (0.1 M). For administration, mice received a single intraperitoneal injection (100 mg/kg) two weeks post irradiation. Following the completion of behavioural testing (i.e., either 4 or 16 days after EdU administration), brains from irradiated and control groups were extracted. We then performed Click chemistry to quantify the number of EdU^+^ cells to assess the efficiency of irradiation on ablation of neurogenesis^56^. For the Click chemistry processing, 50 µm-thick tissues were incubated in Tris buffer of saline (TBS) for 30 minutes. Then the Click solution was prepared by mixing dH_2_O, Tris buffer (1 M, pH = 8.5), CuSO_4_-5H_2_O and azide fluorophore (10mM) in a 599:100:200:1 ratio. Finally, the Click solution was activated by addition of 1M ascorbic acid solution to the rest of the mix in a 1:9 ratio. The tissues were incubated in the activated Click solution for 30 minutes and were washed with PBS three times.

### Specific experimental procedures

#### Generalization experiment (Fig. 1)

For these experiments, mice were handled for approximately 2 mins/day for 3 consecutive days and then were fear conditioned on day 4. Either 2 d or 14 d post-conditioning the mice were tested in the conditioned context or a similar context (order counter balanced, tests 24 h apart).

#### Inception experiments (Figs. 2–5)

For 3 consecutive days, mice received approximately 2 mins of handling and 2-5 mins of cable habituation per day. Next, they were removed from DOX diet for 48 h and were exposed to context A to tag the context. 48 hours after tagging, mice were fear conditioned in context C while receiving photostimulation through the fiber implant. Finally, a recall test was performed either 2 d or 14 d post-conditioning in contexts A and B (order counter balanced, tests separated by 24 h).

#### Inception × irradiation experiments

Mice receive irradiation treatment 3 weeks prior to the behavioral experimentation. After this period, mice were handled and habituated to cable connection for 3 consecutive days. Next, they were removed from DOX diet for 48 h at the end of which the mice were exposed to context A for tagging. 48 h after tagging, mice were fear conditioned in context C while receiving photostimulation. After 2 d or 14 d, the mice were tested in context A or B (order counterbalanced, tests 24 h apart).

### Perfusion, tissue processing and confocal imaging

#### Perfusions

At the completion of behavioural testing, mice were perfused transcardially with 40 ml of cold phosphate buffered saline (PBS; 0.1 M) followed by 40 ml of cold paraformaldehyde (4%; PFA). Brains were extracted, post-fixed in PFA overnight and stored in PBS with 0.3% sodium azide. Brains were switched to an anti-freezing solution (30% sucrose solution) until they submerged, and sliced coronally in 50 µm-thick sections in a cryostat (Leica CM1850). The sectioned tissues were transferred to PBS (0.1 M) in well plates and stored for further processing. In the case of long-term storage, the solution was exchanged with a PBS solution that contained NaN3 (0.2%). All tissues were stored at 4 °C.

#### Immunohistochemistry

Staining was carried out on floating sections in well plates on a shaker. The sections were washed with PBS (0.1M) three times (5 mins each) before staining. Sections were incubated in blocking solution (4% normal goat serum and 0.5% Triton X-100 in PBS) for 2 hours followed by incubation in the primary antibody solution for 72 hours at 4 °C. The primary antibody solution consisted of blocking solution concentrations according to vendor recommendations: chicken anti-GFP (1:1000, Aves, Cat# GFP-1020), mouse anti-NeuN (1:1000, Millipore, Cat# MAB377). Next, sections were washed with PBS (0.1M) three times and were incubated in the secondary antibody solution for 24 hours at 4 °C. The secondary solution was made from 0.1% Tween20 and secondary antibody: goat anti-chicken Alexa Fluor 488 (1:500, Invitrogen, Cat# A-11039), goat anti-mouse Alexa Fluor 647 (1:500, Invitrogen, Cat# A-21235). Finally, sections were incubated in a DAPI (4’,6-diamidino-2-phenylindole) solution (1:10000, Sigma-Aldrich) at room temperature followed by three washes with PBS (0.1M). The sections were then washed and mounted onto gel-coated glass slides and coverslipped using PermaFluor mounting medium (ThermoFisher Scientific, cat# TA-030-FM). The slides were dark-stored overnight to allow the mounting medium to dry.

#### Confocal microscopy

We obtained all images using a confocal laser scanning microscope (LSM 710; Zeiss). We acquired images by scanning 5 z-stacks (5 µm steps) per section with a 20X objective and tiling to cover the region of interest. The laser parameters (e.g. laser power, pinhole size, multiplier gain, and any other digital parameters) were kept the same for all the acquired images of each experiment. The imaging parameters were set using a sample from the control group. The location of each image was selected according to markers of interest per mouse brain atlas.

### Cell counting and analysis

#### Cell counting

For cell counts, the number of cells expressing the marker of interest was quantified. Marker-positive cells in the entire cross-section of the region of interest were counted. All images underwent identical thresholding using the same parameters for each experiment. First, the z-stacks were superimposed, background noise filtered and signals thresholded to acquire countable masks. All the processing was done in Fiji software (National Institute of Health). The counts from the replicate images from each animal were combined to create a final average for each mouse.

#### Statistical analysis

The sample size for the experimental groups was determined based on previous publications^27,30,57^. Mice were pseudo-randomly assigned to different groups with roughly equal numbers of male and female mice. The experimenter was blinded to group IDs when quantifying images and behavioral outcomes. Cell counts were normalized to either DAPI or TRAP counts. The DAPI normalization was used when the overall population was the denominator of interest, and the TRAP normalization was used when the engram population was the denominator of interest. In all experiments, data was analyzed using either t-test or 2-way analysis of variance (ANOVA), with repeated measures when appropriate, followed by post-hoc tests (Holm-Sidak). Statistical significance was set to 0.05 and Bonferroni corrections were used for all analyses.

Different mice were used for all experiments. Statistical analysis was carried out using GraphPad Prism (version 8.0.1).

---

**Figure S1.**
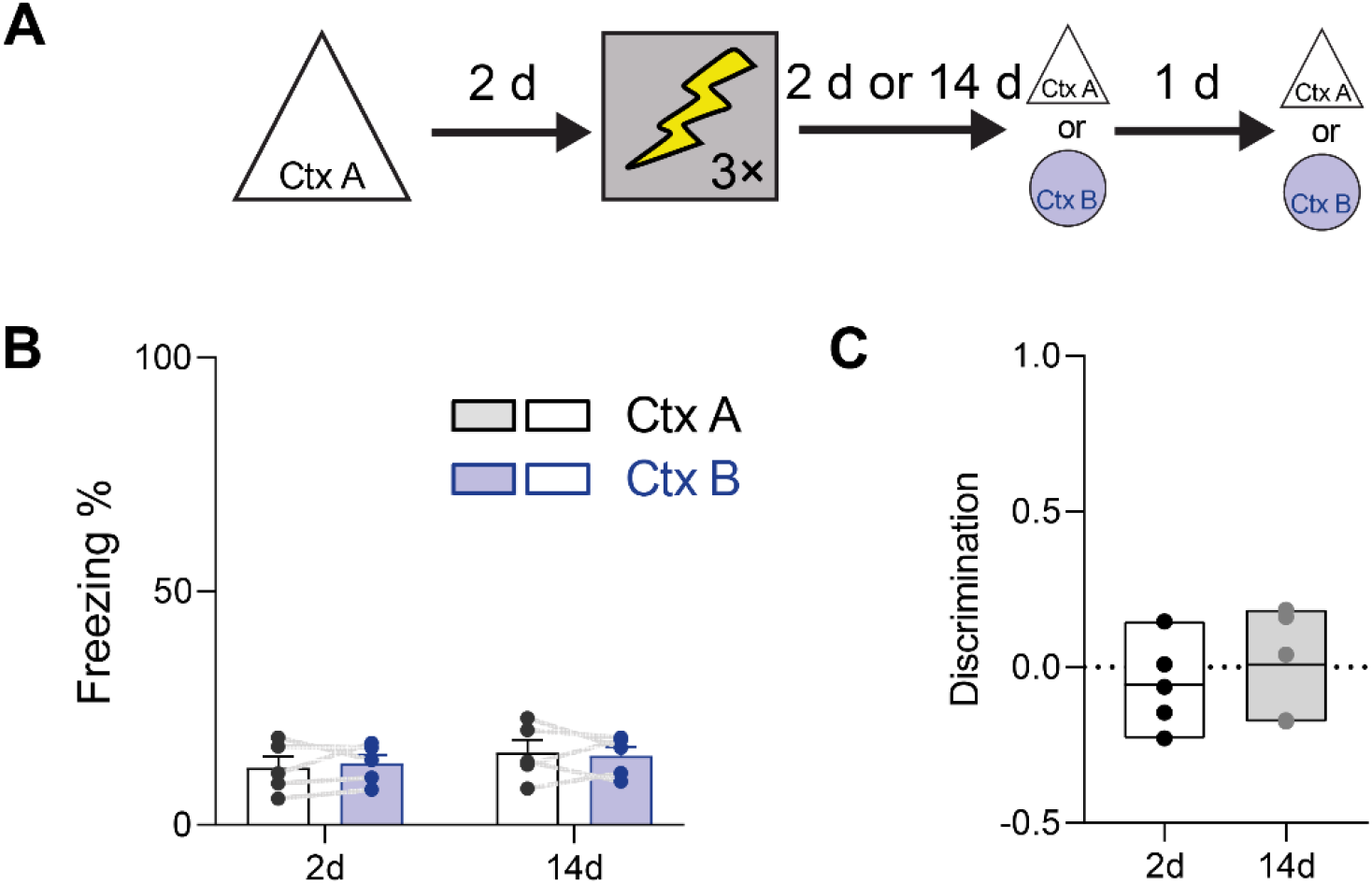
In absence of photostimulation mice do not generalize to contexts A or B. (A) Mice were pre-exposed to context A., and 2 d later trained in context C (in absence of photostimulation). They were then tested in either contexts A or B either 2 or 14 d post-conditioning. (B) Mice did not freeze in either contexts A or B (2-way ANOVA, main context effect: *F*_1,8_ = 0.00, *P* = 0.95; main delay effect: *F*_1,8_ = 0.74, *P* = 0.42; stimulation × delay interaction: *F*_1,8_ = 0.32, *P* = 0.59). (C) No change in the discrimination between A and B at either of the delays was observed (unpaired t-test, t_8_ = 0.64 *P* = 0.54).

**Figure S2.**
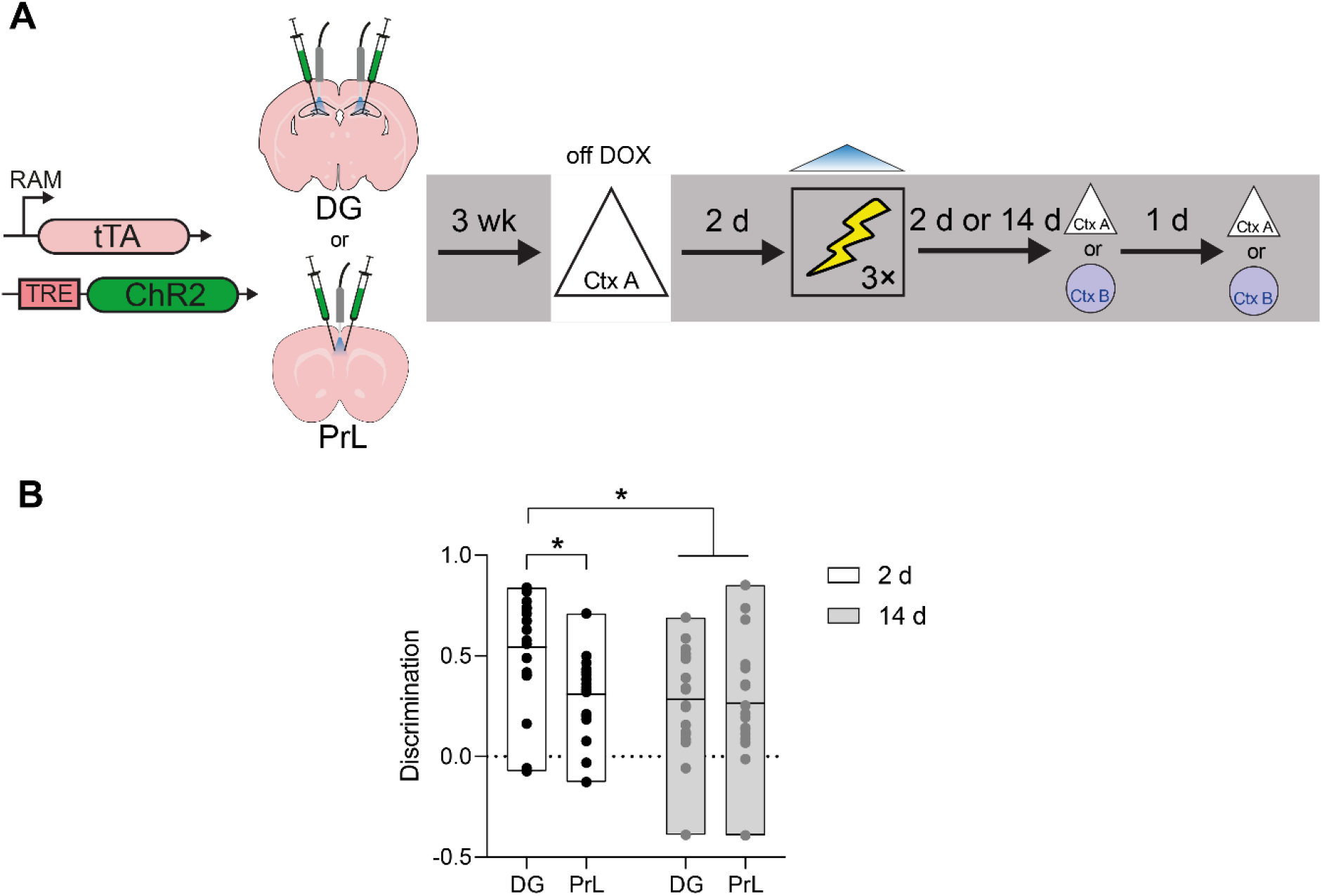
Direct comparison of DG and PrL engram resolution at recent and remote delays. (A) Experimental design (same as Fig. 2 and Fig. 3). (B) Discrimination declines with delay for DG, but not PrL, engrams (2-way ANOVA, main region effect: *F*_1,73_ = 5.72, *P* = 0.02; main delay effect: *F*_1,73_= 5.28, *P* = 0.02; region × delay interaction: *F*_1,73_ = 4.19, *P* = 0.04). * *P* ≤ 0.05, ** *P* ≤ 0.01, *** *P* ≤ 0.001, **** *P* ≤ 0.0001.

